# Epigenetic plasticity is a driver of heritable pollution tolerance in Atlantic killifish

**DOI:** 10.64898/2025.12.18.695277

**Authors:** Mathia Colwell, Melissa K. Drown, Nicole Flack, Carrie Walls, Christopher Faulk, Samantha Carrothers, Emma Weeks, Caren Weinhouse

**Author notes:** Corresponding author: Caren Weinhouse, PhD.

## Abstract

Heritable epigenetic adaptation to environmental stressors is a compelling but highly contested possibility. Previously, we showed evidence of a generationally heritable epigenetic memory at the *cytochrome P450 1a* (*cyp1a*) gene in wild Atlantic killifish (*Fundulus heteroclitus*) with acquired tolerance to polycyclic aromatic hydrocarbons (PAHs). This memory leads to blunted induction of *cyp1a* by PAHs; this blunted response protects against PAH-induced cancer. Here, using Oxford Nanopore long-read sequencing in PAH-tolerant and -sensitive *F. heteroclitus* embryos, we show that PAH-tolerant embryos displayed reduced plasticity in DNA methylation response to PAH, as compared to PAH-sensitive embryos, that was not due to mutational loss of CpG sites. Notably, we observed population differences in DNA methylation of genes in pathways linked to the PAH tolerance phenotype, including aryl hydrocarbon receptor (ahr) and voltage-gated potassium channel signaling, as well as developmental processes and energy metabolism. Specifically, we observed PAH-induced loss of *cyp1a* gene body methylation in PAH-sensitive but not-tolerant embryos. We observed similar patterns at *cyp1b1*, *cyp1c1*, and the aryl hydrocarbon receptor repressor, *ahrr,* which show similarly blunted expression in response to PAH challenge. The reduced loss in genic methylation in tolerant embryos was correlated with greater induction of natural anti-sense RNA transcripts in *cis* (*cis*-NATs), which may regulate transcription of these genes. Our data support the existence of stable epigenetic responses to chronic environmental stressors in a natural experimental setting, with broad implications for natural or directed adaptation strategies for other populations.

## Introduction

Climate change is projected to lead to extreme environmental stress for both animal and human populations^1^. Some animals with short generation times can evolve adaptive phenotypes in response to these stressors^2–7^. However, human populations are long-lived and incapable of rapidly adapting their genotypes to a new environment^8,9^. As a result, there is a strong interest in understanding mechanisms underlying persistent, beneficial acclimation phenotypes in humans in response to environmental stressors^8^. Beyond human-specific outcomes, the relative contributions of genetic adaptation and epigenetic plasticity, and interactions between them, to population responses to rapid environmental change remains a critical area of investigation^10–12^. Like adaptation, acclimation likely brings pleiotropic effects that may be beneficial or adverse, depending on context^10–14^. Therefore, it is critical to understand how these responses function mechanistically to promote beneficial plasticity responses to climate change-related stressors.

Environmental plasticity refers to an organism’s ability to generate more than one phenotype from a given genotype, which confers resilience to a range of environments^8^. Historically, plasticity in physiological or transcriptional states was thought to occur within an individual’s lifetime only and evolutionary adaptations to occur over multiple generations^8,12,15^. However, newer data suggest that epigenetic responses to environmental stressors, which are both a cause and consequence of transcriptional plasticity^9,10^, can lead to generationally persistent adaptations^9,10^. This possibility is highly controversial and data on underlying mechanisms are minimal. In addition, current examples in the literature address environmental stressors that do not have direct translational relevance to human populations. One of the most salient current and future environmental stressors for humans is wildfire smoke^16^, and one of the primary toxic components of wildfire smoke are polycyclic aromatic hydrocarbons (PAHs)^17^. PAHs are common byproducts of incomplete organic combustion that are present in high doses in wildfire smoke, among other forms of air pollution^17^. To address these critical knowledge gaps, we evaluated epigenetic plasticity in response to PAHs in a unique model system, *Fundulus heteroclitus*.

*Fundulus heteroclitus*, colloquially termed mummichog or Atlantic killifish, are a uniquely useful model for studying adaptation to environmental stressors^18^. Mummichog are small teleost fish that live in estuaries along the Atlantic coast^19^ with the ability to survive in a range of extreme environments^18^. Mummichog can tolerate salinity ranging from freshwater to hypersalinity as high as 120%^20^, oxygen concentrations as low as 1 mg/L^21–23^, and a thermal gradient of 12°C^24,25^. In addition to tolerance of natural environmental variation, several specific populations of mummichog have acquired tolerance to extreme levels of local pollution^26^. One of these, located on the Elizabeth River in Virginia, has evolved resistance to the teratogenic effects of extreme exposures to PAHs (reviewed in^27^). This PAH-tolerant phenotype carries documented tradeoffs (reviewed in^27^). PAH-tolerant fish have less robust immune systems and they are more prone to disease^28–33^, they are more susceptible to phototoxicity (a severe acute response to PAH in the presence of UV radiation)^33^, they are less hypoxia tolerant^33^, and they show altered metabolism^34–36^ and reduced thermal plasticity^35–37^.

The PAH-tolerant phenotype and related costs primarily are attributed to rapid genetic evolution in response to extreme selection pressure^26^. Because rapid genetic evolution depends on high standing genetic variation within the affected population, this response has limited value in predicting responses or designing strategies for human populations. However, several lines of evidence suggest that the PAH tolerance phenotype is partially non-genetic, possibly due to generationally persistent epigenetic acclimation responses. First, PAH-tolerant larvae reared in clean water for one to two generations in the laboratory show reduced survival under contaminated conditions and increased survival in clean conditions, as compared to first generation larvae^33^, indicating rapid reversal of PAH tolerance. This result is supported further by a recent report that mummichog from remediated sites on the Elizabeth River are more sensitive to PAH toxicity than are fish from non-remediated sites (although this effect may be due in part to in-migration of wild-type fish)^36^. Second, xenobiotic metabolism genes, including *cytochrome P4501a*, *cytochrome P4501b1*, and *cytochrome P4501c1* (*cyp1a, cyp1b1, cyp1c1*) that are induced by PAH in sensitive fish but less so in tolerant fish recover inducibility when reared in clean conditions^38–40^. Together, these findings are consistent with trait reversion that is too rapid to be explained by genetic change.

Our objective in this study was to address two main questions. First, to what extent does heritable epigenetic acclimation (“epigenetic adaptation”), rather than genetic adaptation, explain the PAH-tolerant phenotype? Second, does reduced epigenetic plasticity explain the costs of the PAH-tolerant phenotype? To answer these questions, we performed Oxford Nanopore long-read sequencing in PAH-tolerant and PAH-sensitive embryos to capture simultaneously genomic sequence and DNA methylation, both at baseline and in response to PAH challenge. We integrated these datasets with paired transcriptomes generated using a short-read, stranded Illumina RNA-seq platform. We observed reduced epigenetic plasticity in response to PAH challenge, as well as differential DNA methylation in genes known to contribute to the PAH-tolerant phenotype and its related costs. Our data support the hypothesis that epigenetic memories of environmental stressors can be generationally inherited and that these responses carry reversible costs. Taken together, these findings have substantial implications for predicting epigenetic plasticity in multiple species, including humans, with increasing environmental stressor exposures.

## Results

### PAH-tolerant fish show reduced DNA methylation response to PAH

To evaluate the extent to which epigenetic plasticity explains the PAH-tolerant phenotype in Elizabeth River *F. heteroclitus*, we compared DNA methylation patterns in F_1_ embryos bred from wild-caught fish from both PAH-contaminated (PAH-tolerant population) and clean reference (PAH-sensitive population) sites on the river. We exposed embryos, rather than adult fish, to avoid experimental bias. PAH-tolerant fish survive poorly in clean laboratory tanks and PAH exposures accumulate over time in fish fat depots^27,40^; therefore, adult fish raised in clean laboratory conditions would be subject to survivor bias and adult fish raised in contaminated conditions would be subject to confounding by pre-existing PAH body burden. In addition, a substantial body of work has been generated in PAH-tolerant embryos, rather than adult fish, to answer questions focused on developmental plasticity. We follow a similar exposure paradigm here, to maximize comparison of our results with the broader field. Briefly, we exposed pools of manually fertilized embryos (*N*=10 pooled embryos per population per condition, *N*=4 total pools) from depurated, wild-caught, F_0_ adult fish to PAH-rich Elizabeth River sediment extract (ERSE), or to clean water, for 14 days post-fertilization. We then performed long-read Oxford Nanopore sequencing of sample genomes and methylomes and short-read Illumina sequencing of sample transcriptomes.

In comparisons between PAH-exposed and control embryos in both populations, most differentially methylated regions (DMRs; defined as at least 1000 bp region containing at least 10 CpG sits with at least 25% difference in DNA methylation) were in genic regions, including exons and introns, and proximal regulatory regions, including 5’ and 3’ untranslated regions (UTRs) (**Fig. 1a**). This result is consistent with a gene regulatory role for DNA methylation in mummichog. In addition, this result supports a role for DNA methylation in environmental plasticity. Previous studies have reported that environmental plasticity responses occur primarily in gene body methylation (GBM)^41,42^. Specifically, these studies show that GBM follows a bimodal distribution, with constitutively expressed genes and inducible genes each clustering near one of the two modes^41,42^. In response to environmental stress, lowly methylated genes gain methylation, and highly methylated genes lose methylation, reducing the distance between modes and ostensibly reducing plasticity to additional stressors^41,42^. Therefore, we hypothesized that the GBM distribution in tolerant fish would be bimodal and would show a narrowed distance between modes, as compared to the GBM distribution in sensitive fish. First, we investigated the GBM distribution in reference datasets (*Homo sapiens, Danio rerio*; **Supp. Fig. 1a**), to provide a reference standard for our experimental results. However, we did not observe a bimodal distribution in GBM in either species (**Supp. Fig. 1a**). We were able to replicate the reported bimodal distribution only when including all genome-wide DNA methylation, not restricted to gene bodies (**Supp. Fig. 1b**). We observed a similar pattern in both mummichog populations in GBM (**Supp. Fig. 1c**) and genome-wide DNA methylation (**Supp. Fig. 1d**). Therefore, our results do not support an increase in GBM uniformity as a mechanism for reduced environmental plasticity.

**Figure 1.**
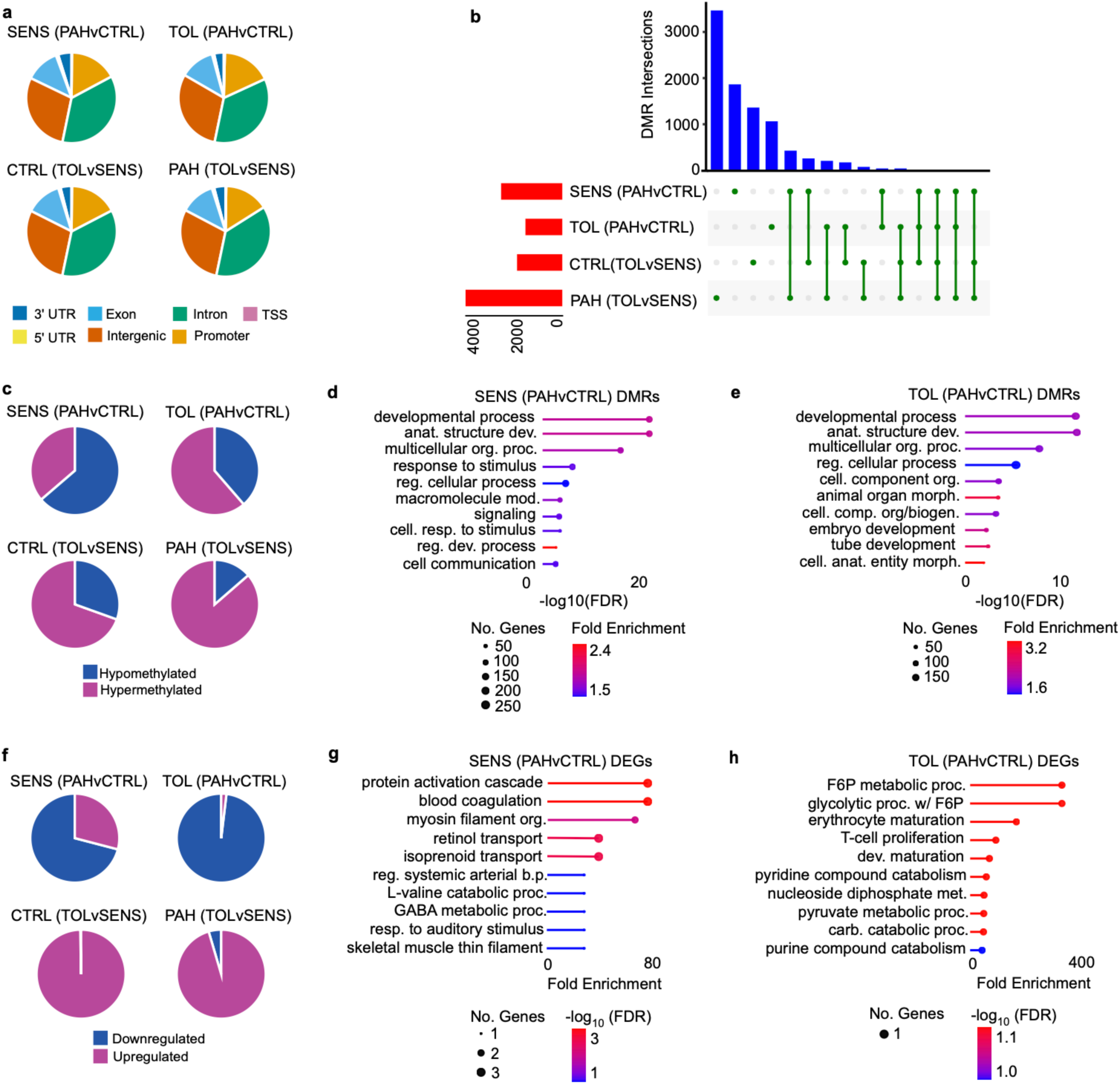
Differential DNA methylation and gene expression in PAH-sensitive and PAH-tolerant *Fundulus heteroclitus*. (a) Genetic context for differentially methylated regions (DMRs) in PAH-sensitive and PAH-tolerant fish, with and without PAH challenge. (b) Upset plot of unique and shared DMRs in PAH-sensitive and PAH-tolerant fish, with and without PAH challenge. (c) Pie-charts of hypomethylated and hypermethylated DMRs in PAH-sensitive and PAH-tolerant fish, with and without PAH challenge. (d-e) Gene Ontology enrichment (top ten terms) in DMRs in PAH-sensitive and PAH-tolerant fish, with and without PAH challenge. (f) Pie-charts of upregulated and downregulated differentially expressed genes (DEGs) in PAH-sensitive and PAH-tolerant fish, with and without PAH challenge. (g-h) Gene Ontology enrichment (top ten terms) in DEGs in PAH-sensitive and PAH-tolerant fish, with and without PAH challenge.

Next, we compared the number of DMRs between populations. We observed substantially fewer DMRs in PAH-tolerant fish (**Fig. 1b**), as compared to sensitive fish (**Fig. 1b**), which is consistent with reduced environmental plasticity. However, there was minimal overlap in DMRs between the two populations (**Fig. 1b**), indicating that tolerant fish are not mounting a reduced response, but rather an alternative one. Sensitive DMRs primarily lost methylation (**Fig. 1c**). In contrast, tolerant DMRs primarily gained methylation (**Fig. 1c**). We note that tolerant fish showed higher DNA methylation levels at baseline (**Fig. 1c**). However, this result does not explain the population difference in PAH response; PAH challenge further increased methylation levels in tolerant fish (**Fig. 1c**).

To evaluate the biological pathways affected by DNA methylation changes, we performed Gene Ontology (GO) enrichment on DMRs from each population (**Fig. 1d-e**). The top hit in both tolerant and sensitive fish was “GO:0032502 developmental process” (**Fig. 1d-e**) which is consistent with the known developmental toxicity of PAHs (reviewed in^18,27^). We note that, of the “top ten” GO terms in each population, four were enriched in both population comparisons (**Fig. 1d-e**). This result indicates that the DNA methylation response was enriched within similar pathways in both populations, despite minimal overlap in DMRs at the gene level (**Fig. 1b**).

Given the known genetic differences between populations, we asked whether the difference in DMR response might have a partly genetic basis. Methylated cytosines are prone to spontaneous deamination, resulting in C>T mutations^43^. Therefore, we evaluated whether our DMR results were confounded by mutational loss of CpG sites. To test whether the DMRs unique to sensitive fish contained CpG sites that were mutated in the tolerant population, we overlapped the C>T mutations in CpG sites in unexposed, tolerant fish with the sensitive DMR set (**Supp. Table 1**). We observed a lower CpG mutation rate (9.82E-04) in sensitive DMRs, as compared to a similar overlap with the tolerant DMR set (2.03E-03) (**Supp. Table 1**). In addition, the number of CpG sites per DMR is unchanged in tolerant fish, as compared to sensitive, at baseline or in response to PAH challenge (**Supp. Fig. 2a-c**). Next, we evaluated the overlap between DMRs and structural variants. We hypothesized that new structural variants recruited the endogenous DNA methylation machinery to silence potential repeat-rich retroelements and stabilize the genome. However, fewer than 10% of DMRs overlapped structural variants in either population (**Supp. Table 2**). Overall, these results show that neither single base substitutions nor structural mutations explained the DMR difference between populations.

**Figure 2.**
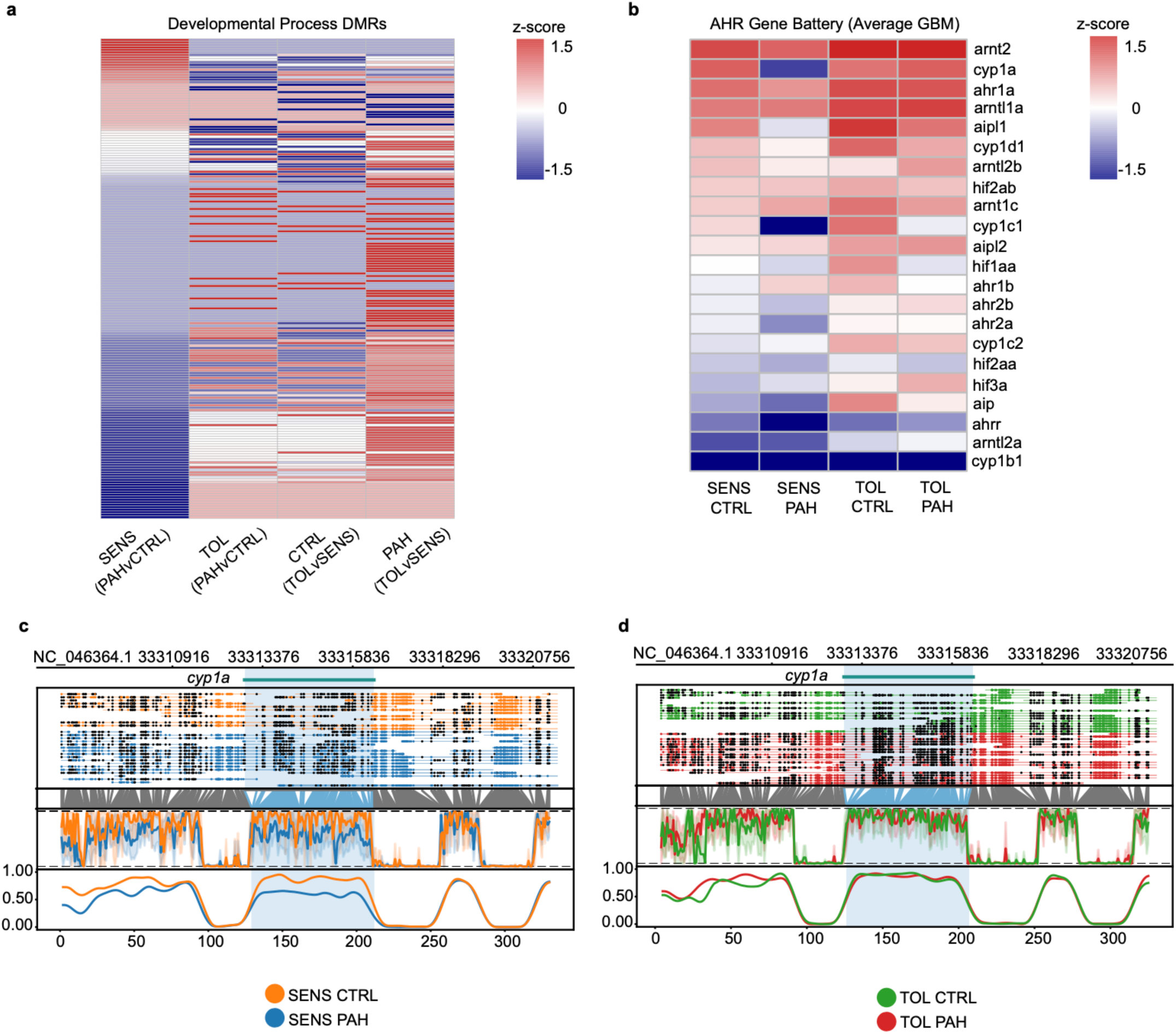
DNA methylation in developmental and ahr battery genes in PAH-sensitive and PAH-tolerant *Fundulus heteroclitus*. (a) DMRs in genes in the Developmental Process Gene Ontology term (GO:0032502) in PAH-sensitive and PAH-tolerant fish, with and without PAH challenge, ordered from largest to smallest effect size in the SENS (PAHvCTRL) comparison. (b) Average gene body methylation (GBM) in the ahr signaling gene battery in PAH-sensitive and PAH-tolerant fish, with and without PAH challenge, ordered from highest to lowest methylation level in SENS CTRL fish. (c) Methylartist plot for the *cyp1a* gene in PAH-sensitive fish, with and without PAH challenge. (d) Methylartist plot for the *cyp1a* gene in PAH-sensitive fish, with and without PAH challenge.

In contrast to DMRs, most differentially expressed genes (DEGs) were downregulated in both populations (**Fig. 1e**); however, nearly all tolerant DEGs are downregulated, as compared to approximately two-thirds of sensitive DEGs (**Fig. 1f**). We note that tolerant fish show substantially higher gene expression than sensitive fish at baseline (**Fig. 1f**). As a result, despite PAH-induced downregulation, tolerant fish still show substantially higher gene expression following PAH challenge, as compared to sensitive fish (**Fig. 1f**). Like DMRs, DEGs show minimal overlap between populations (**Supp. Table 3**). Unlike DMRs, GO enrichment shows dissimilar biological responses between populations; we note that tolerant fish show enrichment in developmental, metabolic, and immune processes that are all linked to the PAH tolerance phenotype (**Fig. 1g-h**). In general, DNA methylation level (within genes or across the genome) correlated poorly with gene expression level of annotated genes (**Supp. Fig. 3**). We speculate that gene expression levels may reflect acute cellular responses and that DNA methylation responses may reflect the potential range of gene expression responses to an environmental stimulus.

**Figure 3.**
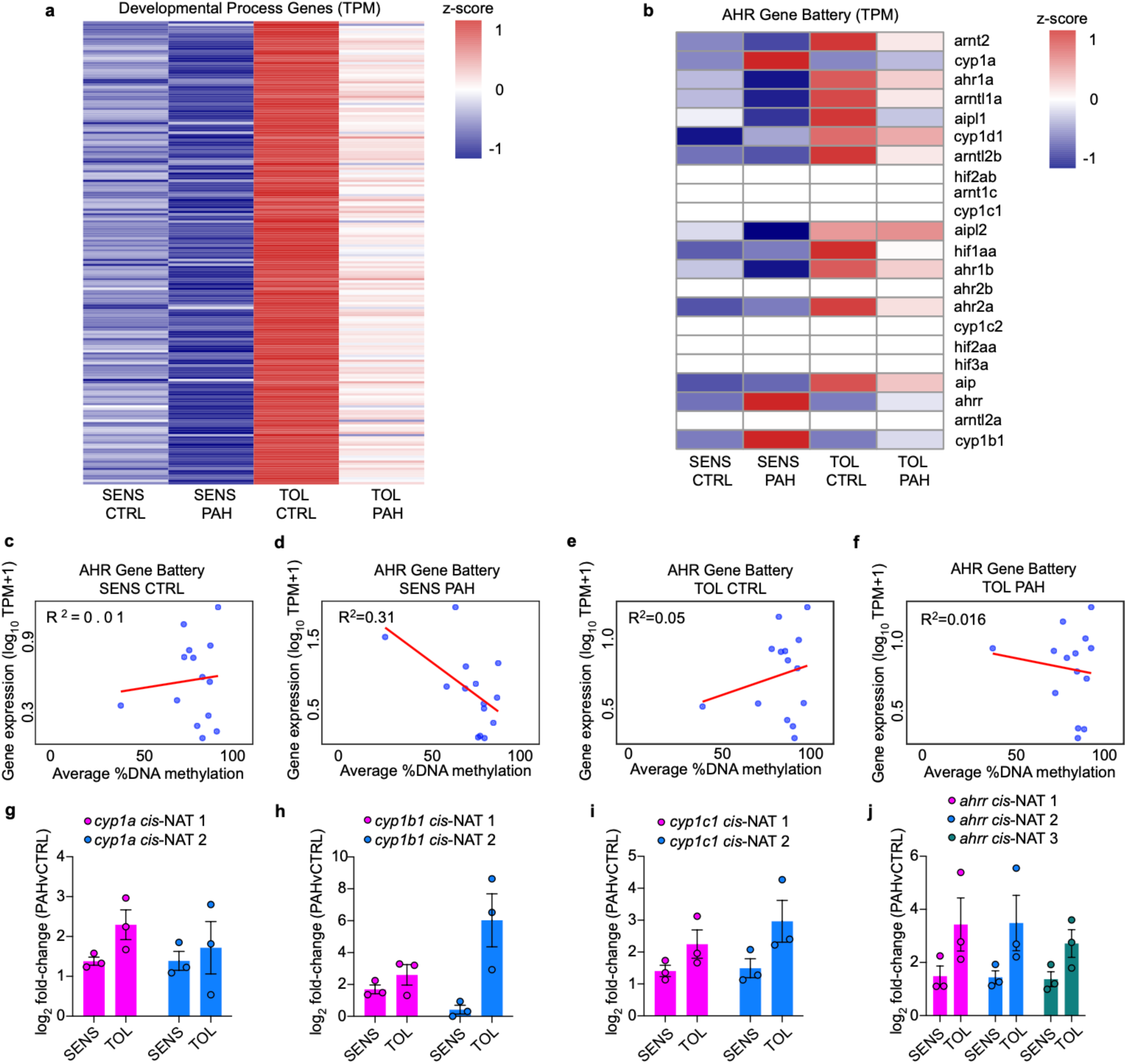
Gene expression in developmental and ahr battery genes in PAH-sensitive and PAH-tolerant *Fundulus heteroclitus*. (a) Expression level for genes in the Developmental Process Gene Ontology term (GO:0032502) in PAH-sensitive and PAH-tolerant fish, with and without PAH challenge, ordered from largest to smallest DMR effect size in the SENS (PAHvCTRL) comparison in Fig. 2a. (b) Expression level for genes in the ahr signaling gene battery in PAH-sensitive and PAH-tolerant fish, with and without PAH challenge, ordered from highest to lowest gene body methylation level in SENS CTRL fish in Fig. 2b. (c-f) Correlation plots for gene expression and DNA methylation for ahr signaling gene battery in PAH-sensitive and PAH-tolerant fish, with and without PAH challenge. (g-j) Bar-plots of gene expression data (generated via qPCR) for candidate *cis*-NATs for *cyp1a*, *cyp1b1*, *cyp1c1*, and *ahrr* in PAH-sensitive and PAH-tolerant fish, with and without PAH challenge.

### PAH tolerance is a partially non-genetic phenotype

Next, we evaluated DMRs enriched in pathways linked to the PAH tolerance phenotype for patterns consistent with epigenetic adaptation to PAHs. We compared DMRs in genes within the “developmental process” GO term (**Fig. 2a, Supp. Table 4**), which was the top hit in both population comparisons. Tolerant fish showed DMR hypermethylation in response to PAH, as compared to similarly challenged sensitive fish (**Fig. 2a, Supp. Table 4**). Many of these DMRs showed higher levels of methylation at baseline in tolerant fish, which was further increased on PAH challenge (**Fig. 2a, Supp. Table 4**). This pattern mimics the overall pattern that we observed in all DMRs between the two populations (**Fig. 1c**). To assess whether similar developmental genes responded to PAH in each population, we evaluated the overlap in the annotated gene sets within the “developmental process” GO term. The DMRs in the sensitive population were annotated by 220 unique genes; the DMRs in the tolerant population were annotated by 135 unique genes (**Supp. Table 4**). Only 41 genes were shared between these two lists (**Supp. Table 4**), and few of the signaling pathways enriched in these genes were shared (**Supp. Table 5**), indicating that the tolerant fish were mounting an alternative developmental program in response to PAH. This result is consistent with a compensatory developmental program contributing to PAH tolerance.

Next, we assessed genes related to aryl hydrocarbon receptor (ahr) signaling. Because many PAHs are ligands for the ahr, we evaluated a battery of genes that encode proteins involved in ahr signaling (including all four *ahr* paralogs), as well as a battery of ahr target genes (including *cyp1a*, *cyp1b1*, *cyp1c1,* and *ahrr*) (**Fig. 2b**). Because methylation differences at these genes did not meet our DMR cutoff of 25%, we compared average GBM for the ahr battery genes (**Fig. 2b, Supp. Table 6**). We observed a 23% drop in *cyp1a* GBM in sensitive fish challenged with PAH (**Fig. 2b-c**), but a gain of 1.5% in exposed tolerant fish (**Fig. 2b, 2d**). We observed a similar pattern at *cyp1b1* (-10% vs. +5%), *cyp1c1* (-26% vs. -11%), and *ahrr* (-9% vs. +2%) (**Fig. 2b, Supp. Fig. 4a-c, Supp. Table 6**).

*F. heteroclitus* have four, well described *ahr* paralogs (*ahr1a/2a*, *ahr1b/2b*), resulting from a tandem duplication of the gene followed by a genome duplication event^44^. *Ahr2a* is linked functionally to the PAH tolerance phenotype^45^. Interestingly, both *ahr1b* and *ahr2b* show an inverted DNA methylation response to PAH in tolerant fish, relative to sensitive (**Fig. 2b, Supp. Table 6**). Both *ahr1a* and *ahr2a* display a similar pattern to the *cyp* genes - a decrease in methylation in sensitive fish that is reduced in tolerant fish (**Fig. 2b, Supp. Table 6**). We note that we did not observe the large *ahr1a/2a* deletion that was previously reported in these PAH-tolerant fish^26^, which, if present, might have contributed to these patterns. In addition, we observed notable patterns at the *ahr-interacting protein* (*aip*) isoforms, including *aip*, *aip-like 1* (*aipl1*) and *aip-like 2* (*aipl2*). Aip is a cytoplasmic binding partner of ahr that controls its nuclear translocation^26^. GBM at all three *aip* isoforms responded to PAH in the same direction in both populations (decreased GBM in *aip* and *aipl1*; increased GBM in *aipl2*) (**Fig. 2b, Supp. Table 6**). However, in each case, tolerant fish showed higher methylation levels both at baseline and on PAH challenge, as compared to sensitive fish (**Fig. 2b, Supp. Table 6**). In summary, our findings support epigenetic regulation of ahr signaling in PAH-tolerant fish.

One of the many functions of ahr signaling is regulation of embryonic development^27^. Aberrant activation of this pathway by exogenous PAHs is thought to be responsible for teratogenesis in PAH-exposed fish^27^. However, developmental toxicity of PAH (specifically, cardiac deformity formation) may be due also to altered function of voltage-gated potassium channels^26,27^. Therefore, we mined our DMR set for genes in this category. We identified 14 DMRs annotated by 12 voltage-gated potassium channel genes (**Supp. Fig. 4a, Supp. Table 7**). In PAH-challenged sensitive fish, 10 DMRs were hypomethylated; PAH-challenged tolerant fish showed the opposite response – 10 DMRs were hypermethylated (**Supp. Fig. 5a, Supp. Table 7**). Despite hypomethylation of eight DMRs in tolerant fish at baseline, when comparing absolute methylation levels in exposed fish from both populations, tolerant fish still showed higher levels at most DMRs in this set (**Supp. Fig. 5a, Supp. Table 7**).

Next, we evaluated gene sets related to established or putative costs of the PAH tolerance phenotype. These include environmentally plastic traits or signaling responses to PAHs that may be lost or altered in PAH-tolerant fish, such as hypoxia response, expression of ion channels that respond to salinity and solute carriers that are involved in membrane transport, metabolism (e.g., genes involved in glycolysis, oxidative phosphorylation, the TCA cycle, mitochondrial function), immune function, estrogen signaling, intracellular calcium signaling, and retinoid X receptor (RXR) signaling. First, we searched for DMRs annotated with genes in any of these categories. We identified five DMRs related to RXR signaling (in the *rxraa* gene), one DMR related to intracellular calcium signaling (in the *ryanodine receptor 1b*, *ryr1b*, gene), four DMRs in three interleukin genes, and 13 DMRs in 11 solute carrier genes (**Supp. Table 7**). In all cases, the majority of these DMRs were hypomethylated in response to PAH in PAH-sensitive fish but hypermethylated in similarly challenged PAH-tolerant fish (**Supp. Fig. 5b, Supp. Table 7**). For gene sets that did not contain DMRs, we explored average GBM for genes previously reported as important for these traits. When relevant, we focused on genes known to be mutated or under positive selection in PAH-tolerant fish^26^. We evaluated nine genes involved in hypoxia response (including four *hypoxia inducible factor*, or *hif* genes and five *aryl hydrocarbon receptor nuclear translocator*, or *arnt* genes (**Fig. 2b, Supp. Table 6**); these genes are listed with the ahr gene battery due to crosstalk between the ahr and its binding partner, arnt, also known as hif1b^46^). Of these, two (22%) hypoxia genes (*arntl2a* and *hif3a*) showed an inverted response in tolerant (hypermethylated) vs. sensitive (hypomethylated) fish challenged with PAH (**Fig. 2b, Supp. Table 6**). In addition, we evaluated nine ion channel genes that regulate salinity acclimation response, 30 genes in estrogen signaling, which crosstalks with ahr signaling, and 49 genes involved in bioenergetic metabolism, which has been reported to be impaired in PAH-tolerant fish (reviewed in^18^) (**Supp. Fig. 5c-e, Supp. Table 8**). In all these gene sets, most genes showed the same inverted pattern (hypomethylation in sensitive fish but hypermethylation in tolerant fish in response to exposure) (**Supp. Fig. 5c-e, Supp. Table 8**).

### Functional outcomes of DNA methylation changes are context specific

To test whether DMRs and GBM differences are functionally relevant, we assessed whether DNA methylation level was associated with gene expression level measured in the same experimental samples. First, we evaluated the full set of DMRs in each PAH-challenged population. In neither was DMR DNA methylation level strongly correlated with expression of DMR-associated genes (SENS R^2^=0.06, TOL R^2^=0.06; **Supp. Fig. 3**). We observed a similar result between gene expression and average GBM across all genes in the genome (R^2^=0.05 in all samples; **Supp. Fig. 3**) and between gene expression and developmental DMR methylation (R^2^=0.01-0.03 in all comparisons; **Fig. 3a, Supp. Table 4**). However, genes in the ahr battery showed stronger relationships with gene expression; in particular, expression of these genes, including expression of several novel splice isoforms, was inversely correlated with methylation in PAH-challenged sensitive fish (SENS PAH R^2^ =0.31; **Fig. 3b-f, Supp. Table 6**). DMR methylation explained a substantial amount of expression variance voltage-gated potassium genes and interleukin genes in the PAH-sensitive population only (voltage-gated potassium channels SENS (PAHvCTRL) DMRs R^2^=0.64, interleukins SENS (PAHvCTRL) DMRs R^2^=0.87) (**Supp. Table 7**). We observed a similar result in GBM in ion channel genes in PAH-sensitive fish at baseline (ion channels GBM SENS CTRL R^2^=0.28) (**Supp. Table 8**). For all other samples/comparisons in all gene sets tested, R^2^ values were less than 0.2. We speculate that DNA methylation level at a given gene or DMR reflects either active or potential gene expression response, depending on the regulatory mechanism governing expression of that gene or gene set. For example, developmental DMRs may represent potential expression of alternative developmental programs that were active earlier in embryonic development than 14 days post-fertilization. In contrast, GBM at ahr genes, particularly *cyp1a*, *cyp1b1*, *cyp1c1* and *ahrr*, may respond to gene induction in real time.

### Blunted induction of AHR genes is associated with increased anti-sense transcription

Last, we evaluated a possible explanation for the association between PAH-induced GBM loss and induction of *cyp1a*, *cyp1b1*, *cyp1c1*, and *ahrr*. A putative function of GBM is to suppress anti-sense transcription^41,42^. Therefore, it follows that GBM loss is permissive of anti-sense transcription. Anti-sense transcripts can repress or induce expression of target genes^47–49^. We hypothesized that GBM loss at these four genes in sensitive fish permitted transcription of anti-sense RNA that augmented induction. Further, we hypothesized that stable GBM at these four genes in tolerant fish repressed formation of these transcripts and blunted gene induction. We identified candidate transcripts (also known as natural anti-sense transcripts that function in *cis*, or *cis*-NATs) using StringTie, a bioinformatics tool that reconstructs the transcriptome, including novel transcripts that are not annotated in the reference genome (**Supp. Table 9**). We identified two candidate *cis*-NATs each for *cyp1a*, *cyp1b1*, and *cyp1c1*, and three candidates for *ahrr* (**Supp. Table 9**). Because all candidate *cis*-NATs contained sequence that did not overlap with the parent gene, we were able to validate expression of these candidates via qPCR (**Fig. 3g-j**). Contrary to our hypothesis, every candidate *cis*-NAT showed higher expression in PAH-challenged tolerant fish, as compared to similarly challenged sensitive fish. We speculate that these *cis*-NATs may recruit DNA methyltransferases to gene bodies to deposit DNA methylation and repress gene induction^50^. Regardless, the consistency of this result strongly suggests that *cis*-NATs play a role in these genes’ blunted induction by PAH in tolerant fish.

## Discussion

Here, we show evidence of epigenetic adaptation, or generationally heritable epigenetic acclimation, to extreme PAH pollution in a population of *Fundulus heteroclitus*. In addition, we show that this adaptation reduced epigenetic plasticity in response to PAH. Because environmental variability within an individual’s lifetime is thought necessary for plasticity to be favored^51^, this reduction is expected if the environment is static (e.g., consistently polluted environment)^13,52^. This theoretical prediction has been confirmed in other natural^14,53,54^ and experimental systems^55–59^. However, it is unclear from these studies what proportion of plasticity loss is due to genetic evolution, as compared to recoverable loss of epigenetic control of gene expression. Prior studies report that clean water-housed PAH-tolerant fish show rapid reversal of some aspects of the tolerant phenotype, including decreased survival when exposed to high levels of PAH and recovery of inducibility of several ahr target genes (reviewed in^18,27^). In addition, epigenetic memories governing blunted induction of ahr genes *cyp1a*, *cyp1b1*, *cyp1c1* and *ahrr* persist for at least one generation in clean water^33,38,39^, which supports epigenetic inheritance of this response that fades in subsequent generations. Taken together, these findings support epigenetic adaptation as a driver of the PAH tolerance phenotype in this population of fish, with implications for other species, including humans.

In this study, most DMRs were located within gene bodies. This finding is consistent with the theory that gene body methylation is a common target of epigenetic plasticity^41,42,60,61^. Methylation within exons and introns is higher in highly transcribed genes, possibly due to greater chromatin accessibility to DNA methyltransferase complexes^62^; alternatively, this methylation may have a functional role in suppressing intragenic or anti-sense transcription^63^. An alternative theory is that gene body methylation controls differential transcription between inducible environmental response genes and constitutively expressed housekeeping genes^41^. In either case, gene body methylation accumulation must confer a strong phenotypic benefit to overcome the increased risk of mutagenesis in protein coding regions^43^. It is possible that the increased gene body methylation that we observed in PAH-tolerant embryos reflects (or maintains) high rates of transcription of environmental stress response genes required to tolerate high PAH exposures. It will be informative to test whether PAH-tolerant fish that show partial reversal of the tolerance phenotype lose some of the accumulated gene body methylation.

The differential DNA methylation signatures that we observed likely reflect a combination of genetic, environmental and gene-environment interaction effects. We observed that very few population-specific DMRs were due to CpG loss and that minimal DMRs overlapped structural variants, indicating that the observed epigenetic responses to PAH challenge were not attributable to genetic variation between populations. However, we cannot rule out gene-environment interactions, or cases in which DMRs’ environmental responsiveness depends, to some degree, on underlying genotype. We note that our genetic variant analysis shows substantial *de novo* variation in PAH-tolerant fish relative to sensitive (**Supp. Tables 1-2**). This result raises the possibility that PAH-induced mutations may provide a substrate for evolutionary selection; in this scenario, PAH exposure would represent both a potent stressor exerting selection pressure and a source of *de novo* genetic variation on which that pressure can act. This possibility adds an important caveat to previous conclusions that rapid evolution of PAH tolerance resulted exclusively from sweeps of standing genetic variants in the population^26^.

Our results have implications for acclimation to climate change. In humans, enhanced epigenetic plasticity is thought to compensate for reduced genetic variation, as compared to *F. heteroclitus* and other, relatively short-lived animals with large effective population sizes. Our findings suggest that, like *F. heteroclitus*, humans may mount effective epigenetic responses to environmental stressors, including extreme PAH levels in wildfire smoke. However, these compensatory responses may carry costs, including reduced epigenetic responsiveness to other environmental stressors. Given the likelihood of future, extreme environmental shifts due to climate change, it is critical to improve our prediction of both beneficial and detrimental epigenetic responses to these exposures to prevent human disease and promote conservation of all impacted species.

## Methods

### Experimental Design

We caught wild *Fundulus heteroclitus* from a PAH-tolerant population from the site of former creosote wood treatment facility (Republic Creosoting) in the southern of the Elizabeth River in Virginia and from a PAH-sensitive reference population from King’s Creek, an uncontaminated tributary of the Severn River in Virginia, as previously described^40^. Briefly, we depurated fish for at least 4 weeks prior to breeding in flow-through systems comprising a series of 30–40 L tanks containing 20% artificial sea water (ASW, Instant Ocean, Foster & Smith, Rhinelander, WI, USA) and maintained adult fish at 23–25 °C on a 14:10 light: dark cycle and *ad libitum* pelleted feed (Aquamax Fingerling Starter 300, PMI Nutrition International LLC, Brentwood, MO, USA). We manually spawned females and fertilized eggs *in vitro* with expressed sperm from males in a beaker containing ASW. Embryos were held for one hour after spawning to allow for fertilization, then washed briefly with 0.3% hydrogen peroxide solution. For PAH exposure experiments, we used a previously collected, processed, and characterized sediment extract (Elizabeth River sediment extract, ERSE) from the Atlantic Woods Industries Superfund site, a former creosote wood treatment facility, in the Elizabeth River (VA, USA). This extract is a real-world mixture of water and suspended solids with a total PAH content of 5,073 ng/mL PAH, summed from analyses of 36 different PAH. We exposed half of the embryos in each group to 5% ERSE (diluted in 20% ASW; this dose induces *cyp* genes in both populations but is not lethal to sensitive fish) and the remaining half to clean water only from 24 hours post-fertilization to 14 days post-fertilization, after which we flash froze embryos in liquid nitrogen and stored at -80 °C. All care, reproductive techniques, and rearing techniques were non-invasive. All vertebrate animal experiments were performed in accordance with all relevant guidelines and regulations following a protocol approved by the Duke University Institutional Animal Care and Use Committee (A139-16-06).

### DNA long-read sequencing, basecalling, and gene annotation

We sequenced the genomes and methylomes of four pools of 10 embryos each, one pool per population-exposure condition. Sequencing was performed on R9.4.1. minion flow cells using Oxford Nanopore’s minKNOW software (v6.3.0), on a custom-built sequencing computer with a NVIDIA GeForce 2080Ti graphics coprocessor (GPU) with 4352 CUDA cores and an AMD Ryzen 3900x processor (CPU) with 12 threads, 24 cores, 64 Gb RAM, and a 1 Tb SSD (solid state drive). Raw signal FAST5 output were converted to POD5 using ONT’s pod5-tools, then basecalled using Dorado (v7.8.3) under the super accurate model^64^. Base modifications were called simultaneously using the Dorado flag –modified-bases 5mC_5hmC.

We filtered basecalled reads to retain reads with a minimum quality score ≥10. Reads were mapped to our Kings Creek *de novo* genome assembly (GenBank BioProject PRJNA1381666, data submitted for publication). Briefly, the Kings Creek genome was assembled using Flye (v2.9.2)^65^ with an estimated genome size of 1.3Gb. Resulting contigs were polished using Medaka (v1.10.0)^66^, with the model (r1041_e82_400bps_sup_v5.0.0), specific to the R9.4.1 sequencing chemistry. To improve assembly contiguity, duplicated contigs were removed using Purge Dups (v1.2.6)^67^. Additional curation was performed to remove contigs with >6x and <60x mean coverage. The resulting draft assembly was scaffolded against the current *F. heteroclitus* reference genome (GCF_011125445.2) using RagTag (v2.1.0)^68^. Gaps were closed using TGS-GapCloser (v1.2.1)^69^. Genome completeness was assessed before and after scaffolding using BUSCO with the Actinopterygii_obd10 lineage dataset. Alignments were parsed using samtools (v1.21)^70^ and custom scripts were used to assign chromosome names. Chromosomes in the final genome were renamed to reflect corresponding chromosomes in the current *F. heteroclitus* reference genome (GCF_011125445.2). Samples showed high mapping rates (99.6-99.8% reads), with a final coverage of 11-13X and post-QC Q30% scores of 66-66.87%.

### Variant calling

We identified small variants using Clair3 (v1.0.10)^71^ specific to our ONT model r941_prom_sup_g5014. Variants that passed the internal quality filters specific to the ONT platform (quality ≥30, allele frequency ≥0.2, coverage ≥10) were used for downstream analysis. Positions with multi-allelic or strand-imbalanced data were excluded. Variant statistics were generated using BCFtools (v1.13)^70^ and VCF tools (v0.1.17)^72^. Bedtools (v2.31.1)^73^ was used to intersect SNPs and DMRs. We designed a custom awk script to identify regional specific SNPs. Structural variants were called using Sniffles (v2.0.2)^74^ using default parameters (SV size ≥ 50bps, quality of ≥25, read depth of ≥10, strand bias filters).

### DNA Methylation

Basecalled BAMs were concatenated, then sorted and indexed with Samtools (v1.21)^70^. Reads were filtered for a minimum mapping quality score of ≥10. The Seqkit stats (v.2.8.2)^75^ command was used to quantify reads lost post-filtering. Samples were aligned to our Kings Creek *F. heteroclitus* reference genome using Minimap2 (v2.28-r1209)^76^. Modkit (v0.3.1)^77^ was used to generate a bedMethyl file by aggregating methylation calls for downstream analysis.

### Differential Methylation

Global, regional, and site-specific DNA methylation were calculated using a custom awk script. Methylation data from modkit were read into the R package methylkit (v1.34.0)^78^ to identify regions of differential methylation across groups. DMRs were identified using methylkit’s logistic regression model with overdispersion correction. Pairwise comparisons were performed between treatment groups. DMRs were defined as 1000 bp tiles with ≥ 10 CpG sites per tile, with a methylation difference ≥ 25% and FDR <0.01. DMRs were annotated to our Kings Creek *F. heteroclitus* reference genome. DMR visualizations were generated using methylartist (v1.3.1)^79^.

### RNA-seq Library Preparation and Sequencing

Raw FASTQ files were processed using FastQC and MultiQC. Across samples, sequencing produced 160-190M paired reads per library. Post-adapter trimming Q30% scores exceeded 88% for all samples. RNA-seq reads (FASTQ) were aligned to our Kings Creek *F. heteroclitus* reference genome using STAR (v2.7.10a)^80^, with parameters specific to stranded RNA-seq data. STAR produced 123-150M mapped reads per sample. STAR alignment had high mapping rates (75.9-81.5% uniquely mapped) with 18-24% unmapped reads, indicating high reference compatibility. The alignment provided coordinate-sorted BAM files for downstream assembly and quantification.

### Transcriptome Assembly and Abundance Estimation

To detect novel isoforms, antisense transcripts, and sense (parent) transcripts, we assembled aligned BAM files using StringTie (v3.0.0)^81^. StringTie outputs included transcript structures in GTF format, cDNA sequences, and transcript abundance levels. Abundance files provided normalized transcript counts (FPKM and TPM). For initial filtering, transcripts with a TPM > 1.0 were retained. Pairwise differential expression was evaluated by calculating log_2_ fold changes; transcripts with greater than ±25% fold change were used for downstream analysis.

### Antisense Transcript Detection and cis-NAT Classification

We identified candidate *cis*-NATs as transcripts with strand-opposite orientation with overlapping sequence with protein-coding genes of interest. CCIVR (v0.2.0)^82^ was used to characterize all global *cis*-NAT species. Candidate *cis*-NATs were visually identified using JBrowse 2 (v2.0)^82,83^ and reported in **Supp. Table 9**. Candidate *cis*-NATs in *cyp1a*, *cyp1b1*, *cyp1c1*, and *ahrr* were validated using qPCR.

### GO Term Enrichment

We performed Gene Ontology (GO) term enrichment for DMR and DEG lists using ShinyGO (v0.85.1) (https://bioinformatics.sdstate.edu/go/), removed redundant or overlapping GO terms using Revigo (http://revigo.irb.hr/), and identified enriched pathways using Enrichr (https://maayanlab.cloud/Enrichr/).

### Analysis Pipeline

An annotated markdown detailing the analysis pipeline is available at: https://github.com/mathiacolwell/Epigenetic_plasticity_PAH_tolerance_killifish

## Data Availability

All data are available on the NCBI GEO database (GEO accession numbers GSE313974 and GSE313975).

---

**Supplemental Figure 1.**
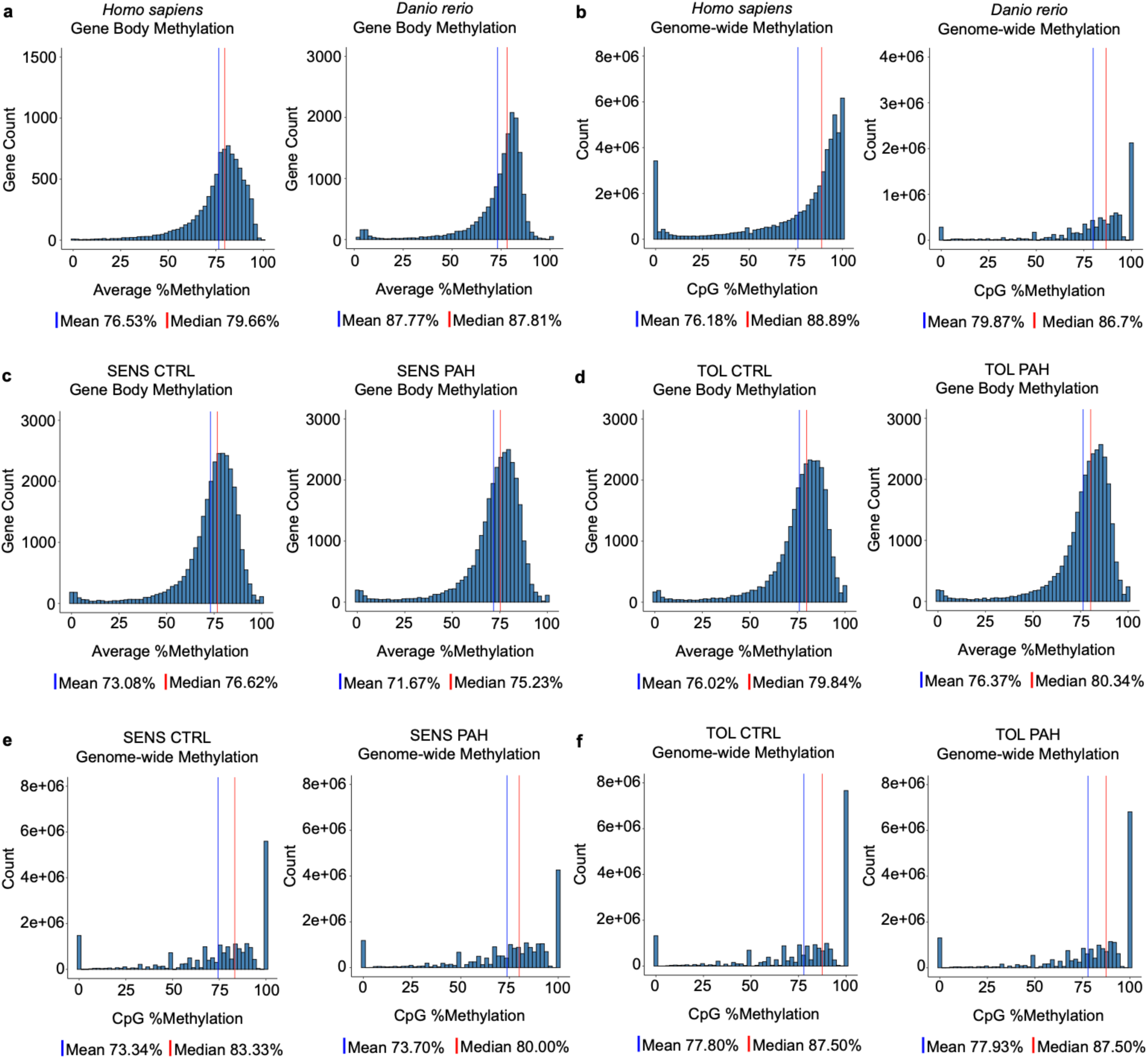
Distributions of gene body DNA methylation and genome-wide DNA methylation in PAH-sensitive and PAH-tolerant *Fundulus heteroclitus*. (a-b) Distributions of gene body methylation (GBM) in reference datasets for humans (*Homo sapiens*) and zebrafish (*Danio rerio*). (c-d) Distributions of genome-wide DNA methylation in reference datasets for human (*Homo sapiens*) and zebrafish (*Danio rerio*). (c-d) Distributions of GBM in PAH-sensitive and PAH-tolerant fish, with and without PAH challenge. (e-f) Distributions of genome-wide DNA methylation in PAH-sensitive and PAH-tolerant fish, with and without PAH challenge.

**Supplemental Figure 2.**
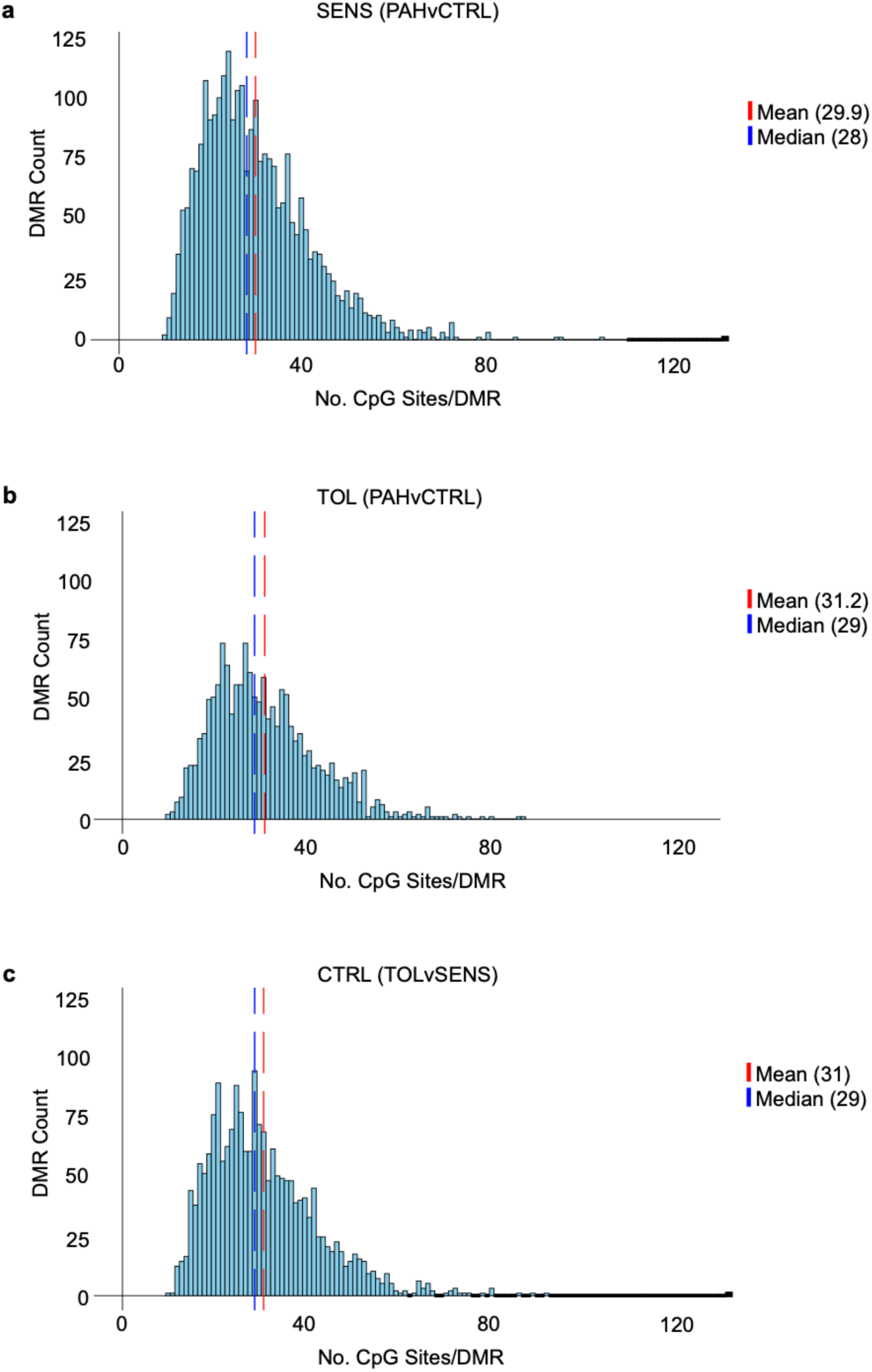
Frequency of CpG sites per differentially methylated region (DMR) in PAH-sensitive and PAH-tolerant *Fundulus heteroclitus.* (a) Histogram of CpG site count per DMR in PAH-sensitive fish (exposed vs. control). (b) Histogram of CpG site count per DMR in PAH-tolerant fish (exposed vs. control). (c) Histogram of CpG site count per DMR in PAH-tolerant fish vs. PAH-sensitive fish at baseline.

**Supplemental Figure 3.**
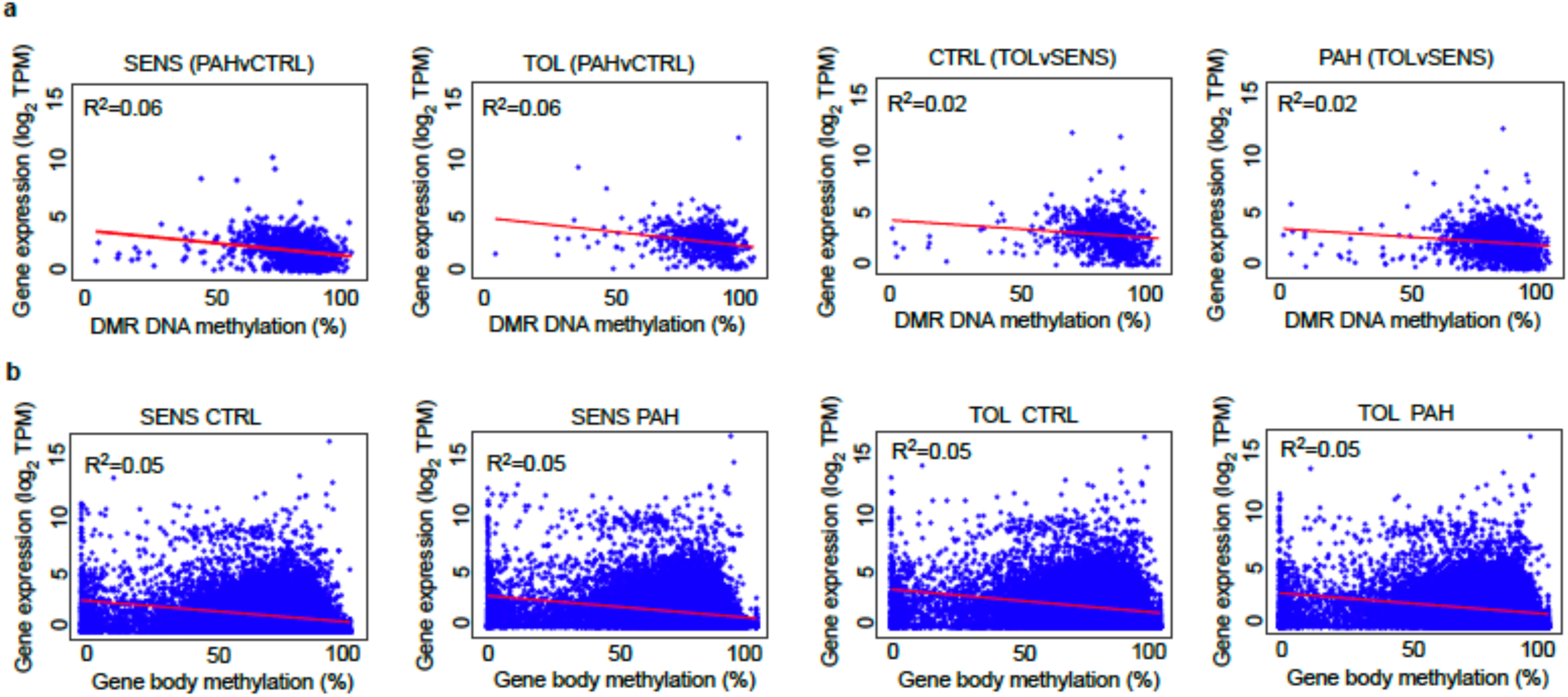
Correlations between DNA methylation and gene expression in PAH-sensitive and PAH-tolerant *Fundulus heteroclitus.* (a) Correlation plots between gene expression and DNA methylation of differentially methylated regions (DMRs) in PAH-sensitive and PAH-tolerant fish. (b) Correlation plots between gene expression and average gene body methylation (GBM) in PAH-sensitive and PAH-tolerant fish.

**Supplemental Figure 4.**
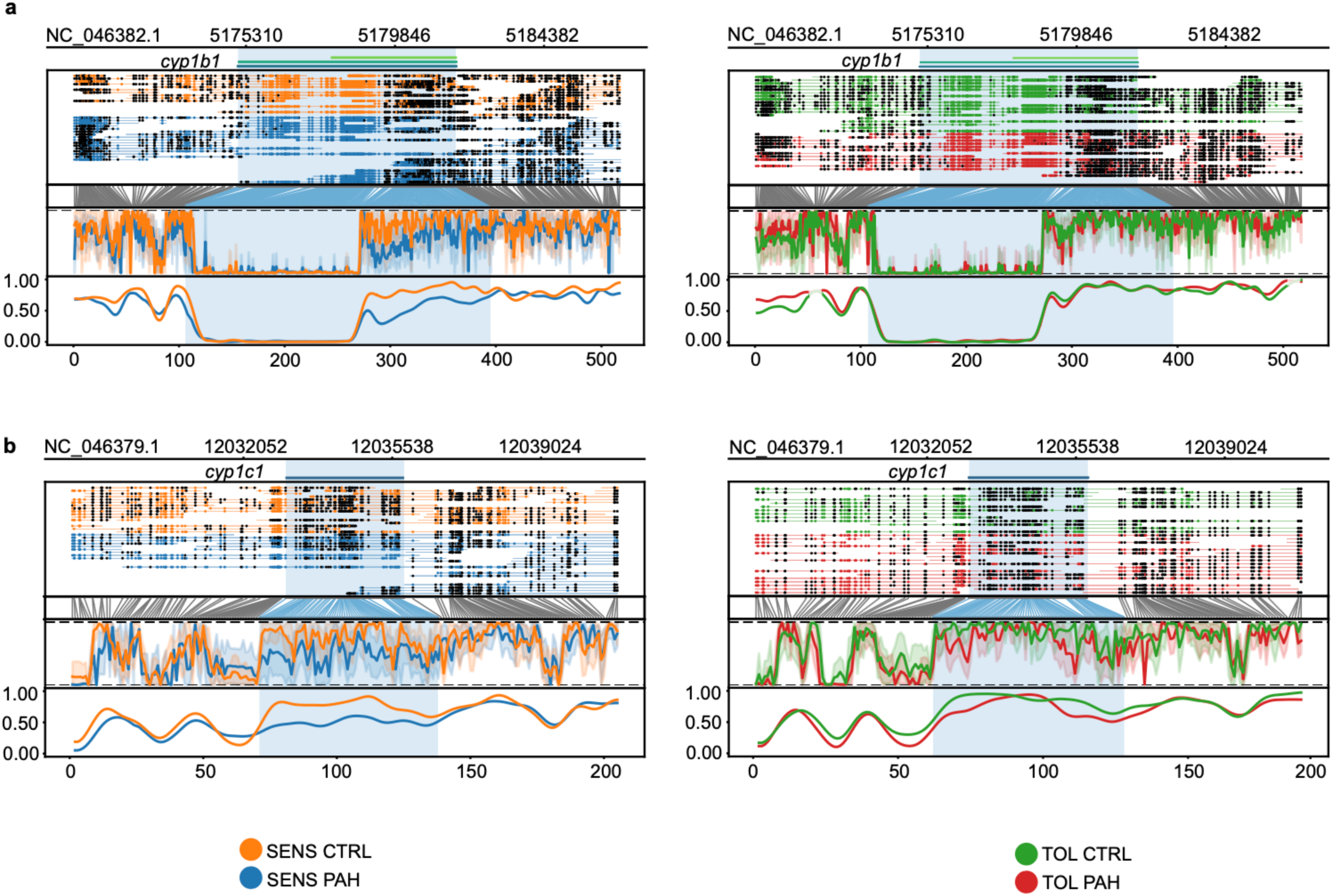
Gene body methylation in ahr battery genes *cyp1b1* and *cyp1c1* in PAH-sensitive and PAH-tolerant *Fundulus heteroclitus*. (a) Methylartist plot for the *cyp1b1* gene in PAH-sensitive fish, with and without PAH challenge. (b) Methylartist plot for the *cyp1c1* gene in PAH-sensitive fish, with and without PAH challenge.

**Supplemental Figure 5.**
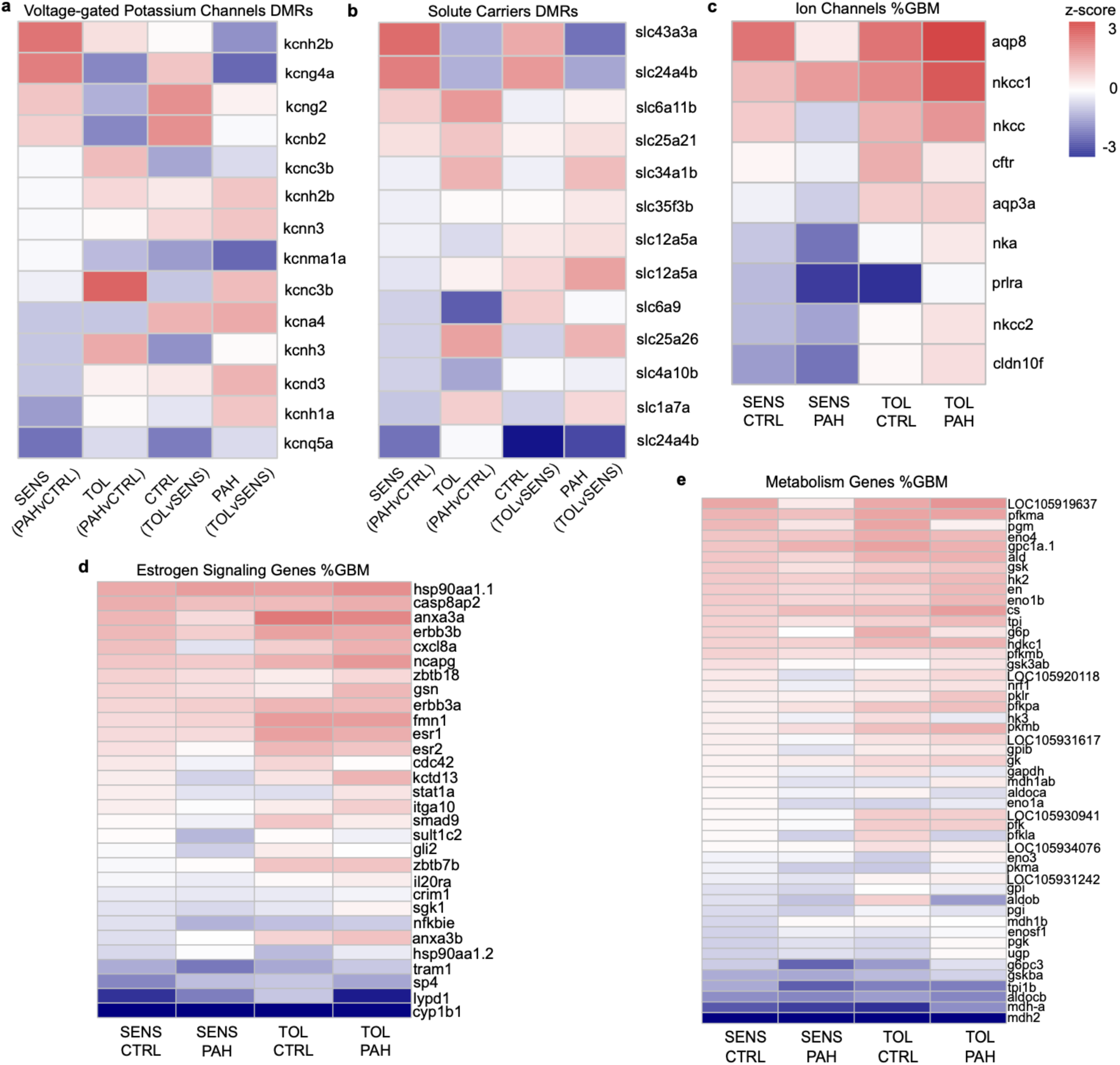
DNA methylation in differentially methylated regions (DMRs) and genes associated with PAH tolerance in *Fundulus heteroclitus*. (a) DMRs in voltage-gated potassium channel genes in PAH-sensitive and PAH-tolerant fish, with and without PAH challenge. (b) DMRs in solute carrier genes in PAH-sensitive and PAH-tolerant fish, with and without PAH challenge. (c) Gene body methylation (GBM) in ion channel genes in PAH-sensitive and PAH-tolerant fish, with and without PAH challenge. (d) Gene body methylation (GBM) in estrogen signaling genes in PAH-sensitive and PAH-tolerant fish, with and without PAH challenge. (e) Gene body methylation (GBM) in metabolism genes in PAH-sensitive and PAH-tolerant fish, with and without PAH challenge. DMR heatmaps are ordered from largest to smallest DMR in SENS (PAHvCTRL). GBM heatmaps are ordered from highest to lowest GBM in SENS CTRL.

